# Calculation of a distribution free estimate of effect size and confidence intervals using VBA/Excel

**DOI:** 10.1101/073999

**Authors:** Joachim Goedhart

## Abstract

Reporting effect sizes aids the transparent presentation and independent interpretation of scientific data. However, calculation and reporting of effect sizes for data obtained in basic research is rare. A standardized effect size was reported by Norman Cliff, known as Cliff's delta. It has several advantageous features, as (i) it makes no assumption on the shape of the underlying distribution, (ii) it works well for small to moderate samples (n>10), (iii) it is easy to calculate, and (iv) its basis is readily understood by non statisticians. Here, a VBA macro, implemented in Excel, is presented. The macro takes two independent samples as input and calculates Cliff's delta with 95% confidence intervals. The macro will reduce the barrier for calculating the effect size and can be a valuable tool for research and teaching.

## Introduction

The use of Null Hypothesis Significance Testing (NHST) for evaluation of scientific data has been highly debated (Goodman, 2008; Cumming, 2014; Nuzzo, 2014). Several papers have highlighted misinterpretation of NHST and resulting p-values (Goodman, 2008; Halsey *et al.*, 2015; Ivarsson *et al.*, 2015; Wasserstein and Lazar, 2016) and have called for use of estimation statistics as alternative (Nakagawa and Cuthill, 2007; Cumming, 2014; Claridge-Chang and Assam, 2016) or additional (Drummond and Tom, 2012; Sullivan and Feinn, 2012) strategy for data analysis and presentation.

Here, I only treat the case in which the data is obtained from a randomized experiment on two independent groups. The NHST returns a p-value that indicates the probability that the data from the two groups is identical, i.e. the null hypothesis is true, given the observed data or more extreme values. If the p-value is below a predefined, arbitrary threshold, usually p<0.05, the result is explained as evidence in favor of an alternative hypothesis, with smaller p-values taken as stronger evidence in favor of the alternative hypothesis. Importantly, p-values do not signify the strength of evidence in favor of an alternative hypothesis (Goodman, 2008; Schneider, 2015). Moreover, NHST and the resulting p-value do not give any information on the magnitude of the difference (Nakagawa and Cuthill, 2007; Cumming, 2014; Ivarsson *et al.*, 2015; Motulsky, 2015). To obtain information on the magnitude of the difference or the size of an effect, the effect size needs to be calculated (Sullivan and Feinn, 2012; Cumming, 2014). The effect size is arguably the parameter that is of interest, since it is related to the biological (or clinical) phenomenon that is studied (Nakagawa and Cuthill, 2007). Nevertheless, estimation statistics is rarely used in basic research and reporting NHST dominates (Tressoldi *et al.*, 2013).

Correct calculation of effect sizes for data that deviates from the normal distribution is rare. To enable wide utilization of effects sizes in basic research, I draw attention to a standardized effect size known as Cliff's delta, that does not make assumptions on the underlying distribution (Cliff, 1993, 1996; Vargha and Delaney, 2000). The Cliff's delta was originally derived to measure effect size on ordinal data, often encountered in psychology. It works equally well for data consisting of quantitative, continuous variables, which is the predominant output in basic research (Vargha and Delaney, 2000; Hsu, 2004). Of note, Cliff's delta is a linear transformation of the A value reported by Vargha and Delaney (Vargha and Delaney, 2000). Both effect sizes were shown to be particularly robust in case of small to moderate (10-50) sample sizes with a non-normal distribution (Delaney and Vargha, 2002; Feng and Cliff, 2004; Li, 2015).

The calculation of Cliff's delta involves the comparison of all values from dataset A with that of dataset B. When a value from set A is larger than that of set B +1 is noted and in the reverse situation −1 is noted. In case of ties 0 is noted. The comparison of set A and B can be graphically represented in a dominance matrix (Cliff, 1993), see figure 1B for an example. Summing all the noted values and dividing through the total number of counts yields Cliff's delta. Negative value indicates that B dominates over A and positive values show that A dominates B. Cliff's delta can also be calculated from the common Mann-Whitney U statistic (Cliff’s delta = 2U/n_A_n_B_ − 1).

**Figure 1.**
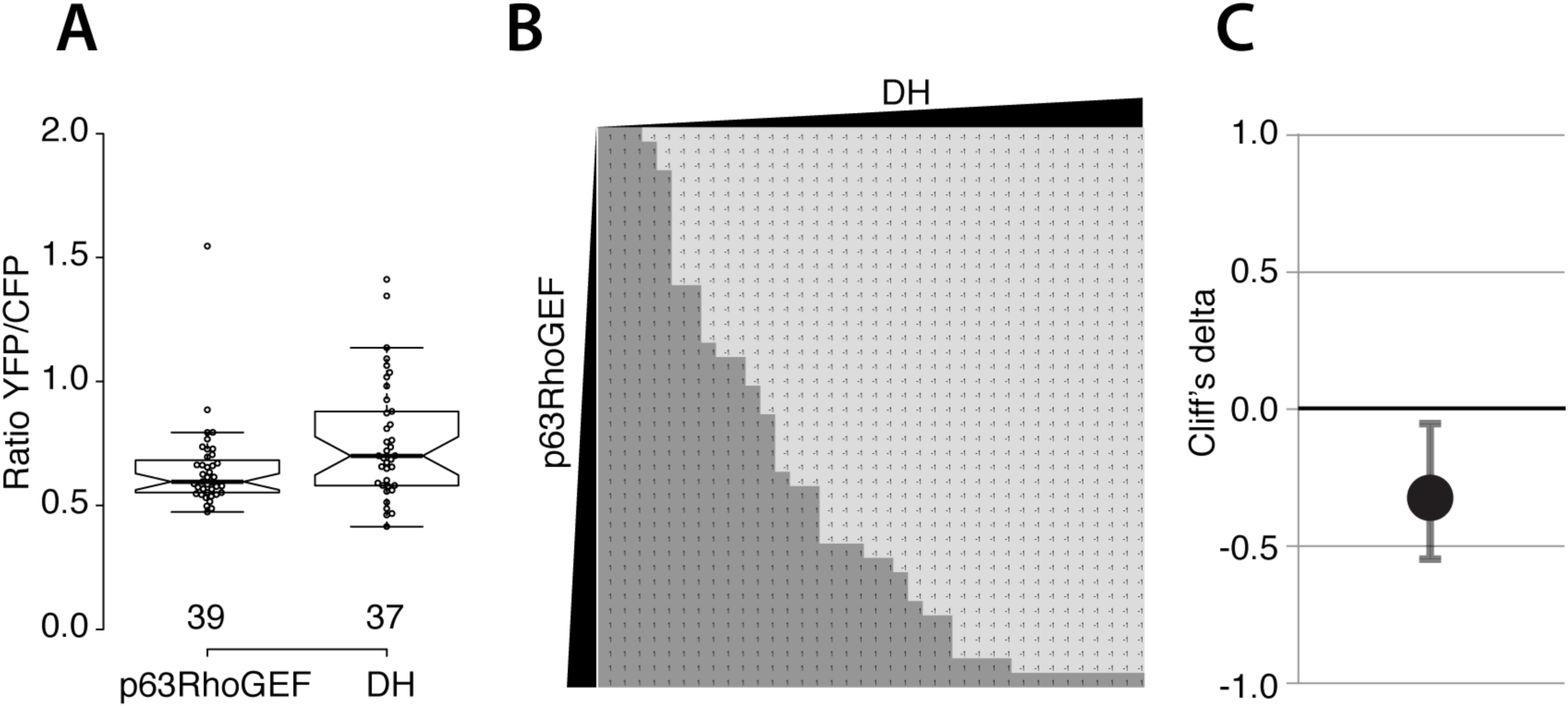
Example of the calculation of the dominance matrix and Cliff’s delta. (**A**) The individual data points shown as open circles indicate the Ratio YFP/CFP, which represents the RhoGTPase activity. The values are shown in a box plot for two conditions, p63RhoGEF and DH. The centerlines show the medians and the box limits indicate the 25th and 75th percentiles. The notches represent the 95% confidence interval for each median. The whiskers extend 1.5 times the interquartile range from the 25th and 75th percentiles. (**B**) The dominance matrix, which is generated by the macro. All values from a dataset are sorted from low to high (indicated by the black bars with increasing width). The matrix is filled by comparing all the data points from both sets. Light grey indicates −1 corresponding to the case DH>p63RhoGEF and dark grey indicates >+1 when DH<p63RhoGEF. (**C**) A graph that shows the resulting Cliff’s delta (-0.32) and the error bars indicate the 95% confidence interval [−0.05, −0.55].

The absolute value of Cliff's delta ranges from 0, i.e. no effect, to 1.0, indicating a maximal effect. Since the effect is standardized, it is possible to discern different categories. Based on the categories first defined by Cohen, Vargha and Delaney (2000) calculated that Cliff’s d effect sizes of 0.11, 0.28 and 0.43 correspond to small, medium and large effects respectively. These categories may serve as rough guidelines for interpreting effect sizes and should not be taken as strict rules, since the effect size should be interpreted and judged in the full context of the experiment (Nakagawa and Cuthill, 2007; Cumming, 2014). Cliff's delta has several advantageous features (Cliff, 1996; Vargha and Delaney, 2000; Hsu, 2004; Ruscio, 2008) and its most powerful aspect is the straightforward calculation and the intuitive interpretation, which can be aided by a graphical representation of the dominance matrix (Cliff, 1993). Moreover, Cliff's delta (i) needs no assumption on the underlying distribution, (ii) is robust in case of outliers or skewed distributions and performs well for normally distributed data (iii) allows comparison for samples with unequal sample size and (iv) works well for small to moderate sample sizes (n>10).

### Rationale and development of the macro

Several options to calculate Cliff's delta and confidence intervals (CI) can be found on the web. These require specialized statistics packages such as R (http://cran.r-project.org/web/packages/orddom/index.html) or SAS (http://citeseerx.ist.psu.edu/viewdoc/summary?doi=10.1.1.488.2246). To allow for a broader adoption of Cliff's delta, I have developed a visual basic macro that runs in Excel. The macro has been tested and shown to work using Mac OS 10.9.5 with Microsoft Excel 2011 (version 14.6.0) and using Windows 7 with Microsoft Excel 2010 (version 14.0.7163.5000). It takes two datasets as input in column A and B and calculates Cliff's delta and the asymmetric 95% CI around the point estimate using equation 5 (Feng and Cliff 2004). In addition, it presents the dominance matrix on a separate, second sheet. A third sheet presents several parameters that are used to calculate the consistent estimate of the variance and the unbiased estimate of the variance (Cliff, 1993). The 95% CI derived from the two variances are also listed.

## Results

The macro is applied to a dataset previously published on the effect of several protein variants on RhoGTPase activity in single cells and the results are summarized in figure 1. The RhoGTPase activity is measured with a FRET biosensor and yields a YFP/CFP ratio value for individual cells, where a higher ratio correlates with higher activity. In one specific condition, we examined the effect of p63RhoGEF versus a truncated variant, DH in Hek293 cells. The data was first reported by van Unen in supplemental figure S2 (van Unen *et al.*, 2015) and analyzed with NHST resulting in a p-value of 0.015 (two-tailed Mann-Whitney test). Here, the individual data points of that dataset are depicted and a boxplot (Spitzer *et al.*, 2014) is used to summarize the data (figure 1A). It can be inferred from the figure that both datasets have a non-normal distribution and contain some extreme values, arguing against calculation of an effect size that assume a normal distribution. Calculating the Cliff's delta results in a value of −0.32 [−0.05, −0.55]. The effect can also be appreciated by inspection of the dominance matrix (figure 1B), showing that DH values are generally higher that those for p63RhoGEF. A graphical representation of Cliff’s delta and the 95% confidence interval is depicted in figure 1C.

## Conclusion

To conclude, I report on a VBA macro implemented in Excel for calculating Cliff's delta and its 95% confidence interval for a two-group randomized experiment. This tool should lower the barrier for calculating effect sizes in basic research and it can be used for teaching to explain the calculation of Cliff’s delta.

## Acknowledgments

I would like to thank Paul Goedhart (Wageningen UR, The Netherlands) for comments and Marten Postma (University of Amsterdam, The Netherlands) for explaining statistical concepts and enlightening discussions.

## Supplemental Material

Text-macro-160905.txt

VBA macro to calculate Cliff’s delta and CI and dominance matrix.

Excel-CLIFFS_DELTA_160905.xlsm

An Excel workbook with the macro and data presented in figure 1.

